# Eavesdropping roots: *Fagus sylvatica* detects belowground stress signals from conspecific and heterospecific (*Picea abies*) neighbors, triggering increased shoot VOC emissions

**DOI:** 10.1101/2025.10.30.685342

**Authors:** Mirjam Meischner, Simon Haberstroh, Jürgen Kreuzwieser, Jörg-Peter Schnitzler, Christiane Werner

## Abstract

- Volatile organic compounds (VOCs) facilitate aboveground plant communication, but belowground signaling remains less understood.
- This study explored root-root interactions between *Picea abies* and *Fagus sylvatica* saplings in monospecific (*Fagus-Fagus*) and heterospecific (*Picea-Fagus*) pairs (n=6), excluding shoot-level VOC communication. Sender plants were treated with jasmonic acid to simulate herbivory and labeled with ^13^CO_2_ and ^15^NH_4_NO_3_ to trace nutrient transfer in a split-root design. VOC emissions and gas exchange were measured over ten days using PTR-TOF-MS and ^13^CO_2_-spectroscopy and ^13^C and ^15^N were analyzed in roots and shoots via EA-IRMS.
- Our findings reveal, that (i) JA treatment induced strong *de novo* terpenoid emissions from *P. abies* and enhanced emissions of oxygenated VOCs and benzenoids from *F. sylvatica*, (ii) *F. sylvatica* receiver plants responded similarly to JA-treated neighbors, indicating belowground signaling, and (iii) responses of receiver plants were more pronounced in the heterospecific treatment. Furthermore, formic acid emissions from soils increased following JA treatment, suggesting altered soil microbial activity. Isotopic analysis revealed C exudation into the rhizosphere and N transfer to receiver plants.
- These results suggest that belowground signaling enables early priming of herbivore-induced defenses in neighboring plants, and that the response intensity is modulated by species identity.

## Introduction

Plants have evolved the capacity to perceive cues from their environment in order to enhance their fitness and adapt to changing environmental conditions (Bigot *et al*., 2018; Novoplansky, 2019; Karban, 2021). They can detect neighboring plants based on the ratio of red to far-red light (Casal *et al*., 1990; Novoplansky *et al*., 1990), adjust their root growth towards favorable nutrient sources (Drew & Saker, 1975; Jackson & Caldwell, 1996) and water availability (Takahashi, 1994; Dietrich, 2018), and respond to mechanical stimuli (Biddington, 1986; Braam, 2005). Plants also produce and are sensitive to chemical cues, especially volatile organic compounds (VOCs). These play a crucial role in their interactions with the environment and other organisms (Kesselmeier & Staudt, 1999; Kessler & Baldwin, 2001; Loreto *et al*., 2014). VOCs are a large and diverse group of small, lipophilic compounds with high vapor pressure (Pichersky *et al*., 2006; Dudareva *et al*., 2013), which are produced by plants to increase their level of resistance against abiotic and biotic stressors, such as drought, heat, and pathogens (Karban & Baldwin, 2007; Loreto & Schnitzler, 2010; Possell & Loreto, 2013). Since the first evidence for VOC-mediated signaling in the early 1980s (Rhoades, 1983; Baldwin & Schultz, 1983), the role of VOCs in plant-insect (Dicke *et al*., 1993; De Moraes *et al*., 1998; Clavijo McCormick *et al*., 2012), within-plant (Frost *et al*., 2007; Heil & Silva Bueno, 2007; Li & Blande, 2017) and aboveground plant-plant signaling (Kessler *et al*., 2006, 2023; Baldwin *et al*., 2006) has been extensively studied. This revealed complex ecological relationships between induced VOC emissions, herbivore behavior (attraction/repulsion) and physiological responses in neighboring plants, such as priming of defensive genes (Kessler *et al*., 2006; Frost *et al*., 2008). It is noteworthy that VOCs released by sender plants can exert a dual effect on the defensive traits of receiver plants, exhibiting positive (Karban *et al*., 2014) and negative (Erb, 2018) outcomes. It has been demonstrated that they, for example, induce the accumulation of proteinase inhibitors in receiver plants, thereby increasing their resistance to pathogen attacks (Farmer & Ryan, 1990). Alternatively, they can increase the susceptibility of receiver plants to herbivores resulting in enhanced leaf damage (Li & Blande, 2015). Another frequent response to VOC exposure in receiver plants is an increase in VOC emissions, such as green-leaf-volatiles (Engelberth *et al*., 2004) and isoprenoids (Ruther & Kleier, 2005). This response has been shown as a defense mechanism potentially mitigating the damage by leaf herbivory (reviewed by Brosset & Blande, 2022). In view of accelerating climate change and the consequent enhancement of the frequency and severity of insect outbreaks in the future (Bale et al., 2002; Pureswaran et al., 2018; Harvey et al., 2020), there is pressing need for more profound understanding of the plants’ defense strategies, such as signal propagation to neighboring plants.

While the subject of aboveground signaling via VOCs has been extensively studied, comparatively little research has been conducted on belowground signaling, which involves a complex interplay of volatile and non-volatile cues (Mathieu *et al*., 2024). The latter include, but are not limited to, root exudates, which play a crucial role in mediating interactions between roots and their environment (Wang *et al*., 2021; Yoneyama & Bennett, 2024), shaping microbial communities (Hu *et al*., 2018; Zhou *et al*., 2023) and facilitating neighbor recognition (Semchenko *et al*., 2014; Kong *et al*., 2018). Furthermore, the mediation of belowground interactions by mycorrhizal fungi has been demonstrated to occur through the transfer of nutrients (Simard *et al*., 1997; Egerton-Warburton *et al*., 2007) and the transfer of signals regarding herbivore and pathogen attacks (Song *et al*., 2010; Babikova *et al*., 2013b; Gilbert & Johnson, 2017). Despite the evidence for resource transfer among plants, the quantitative contribution of specific transfer pathways, such as transfer via common mycorrhizal networks or uptake from root exudates dissolved in the soil solution, remains a subject of debate (Hoeksema, 2015; Karst *et al*., 2023; Klein *et al*., 2023). While the transfer of such resources through biological systems can be studied by stable isotope labeling of the respective compounds, such as nitrogen (^15^N) or carbon (^13^C), the analysis of belowground VOCs is rather challenging due to methodological constraints. Nevertheless, a diverse blend of alcohols, oxygenated compounds, organic acids and terpenoids have been shown to be released from roots (Steeghs *et al*., 2004; Meischner *et al*., 2024), and the activity of terpene synthases (TPS) has been demonstrated in roots of several species, e.g. *Arabidopsis thaliana* and *Centaurea stoebe* (Vaughan, 2010; Gfeller *et al*., 2019). Although root VOCs are partially degraded by soil microorganisms (Rinnan & Albers, 2020; Honeker *et al*., 2023; Pugliese *et al*., 2023; Jiao *et al*., 2023), specific compounds, such as (E)-β-caryophyllene and pregeijerene, are involved in tritrophic interactions between herbivorous insects, their host plants and natural enemies attracted by induced root VOCs (Rasmann *et al*., 2005; Rasmann & Turlings, 2008; Ali *et al*., 2011; Hiltpold *et al*., 2011). Similar to aboveground plant-plant signaling, root VOCs can exert both negative and positive effects on neighboring plants. For instance, they may possess allelopathic properties (Ens *et al*., 2009; Jassbi *et al*., 2010), rendering neighboring roots more susceptible to herbivores (Robert *et al*., 2012; Huang *et al*., 2019), or, promoting the germination rate and growth of receivers (Gfeller *et al*., 2019). In a split-root experiment, which allows for the manipulation of specific parts of the root system, while also enabling direct root interactions with neighboring plants, Falik *et al*. (2012) demonstrated that information about drought was transmitted through the root system, leading to stomatal closure in well-watered plants.

We hypothesize that in accordance with the various signal transduction pathways from root to root, root-root signaling can result in changes in VOC emissions from receiver plants growing adjacent to stressed sender plants analogous to aboveground signaling.

In order to investigate this hypothesis, *Picea abies* and *Fagus sylvatica* saplings were planted in pairs in a split- root design. This allowed root contact, but airborne shoot-shoot signaling was experimentally excluded. These two species were selected as representatives of the key tree species in Central European forests with distinct defense strategies: *P. abies* is a coniferous species with specialized VOC storage pools (Ghirardo *et al*., 2010; Niinemets *et al*., 2011), and *F. sylvatica* a deciduous broadleaved species with labile storage pools and generally lower VOC emission rates (Dindorf *et al*., 2006; Holzke *et al*., 2006).

In order to examine the impact of species combination on signal transfer, we investigated the interactions between *P. abies* and *F. sylvatica*, as well as intraspecific interactions between two individuals of *F. sylvatica*. To mimic the effects of herbivory, the shoots of sender plants were treated with a jasmonic acid (JA) solution. This is a well-established method for inducing a defense response that is similar to the effects of herbivore attack (Degenhardt *et al*., 2010; Li *et al*., 2019; Meischner *et al*., 2024). The experimental approach adopted in this study comprised three main steps. First, the nutrient transfer between sender and receiver plants was investigated as an indicator of active root-root interactions by labeling sender plants with ^13^C and ^15^N and subsequently determining isotopic ratios in the plant tissues. The second step involved the characterization of the response of photosynthetic gas exchange and VOC emissions from the sender plant’s shoots to simulated herbivory. These responses were continuously monitored by ^13^CO_2_ spectroscopy and PTR-TOF-MS, respectively. The third step entailed the evaluation of the responses exhibited by the receiver plants. Furthermore, to assess changes in soil biochemistry due to simulated herbivory, we analyzed soil VOC emissions and soil respiration, both directly affected by root activity. By exploring these processes, this study aims to improve our understanding of how plants respond to environmental stressors and whether defense- related traits, such as VOC emissions, undergo change in neighboring plants due to root-root signaling.

## Material and methods

### Plant material and experimental design

Two-year-old saplings of *Fagus sylvatica* L. (European beech) and *Picea abies* L. (Norway spruce) from a local tree nursery (Zell am Harmersbach, Germany) were planted in spring 2021 using a split-root design (Figure 1). A sender plant and a receiver plant were each grown together in a custom-made box (300 x 400 x 320 mm) which was divided into three compartments by two sealed partitions made of acrylic glass and filled with soil substrate (60 vol.% Floradur, 40 vol.% sand, 5 mg L^-1^ NPK fertilizer and 20 mm of foamed clay for water drainage). In this set-up, plants shared a central soil compartment with their root systems, while also rooting to unshared compartments on the outer sides. Either *P. abies* or *F. sylvatica,* were planted on the sender position with *F. sylvatica* on the receiver position, resulting in two species combinations: *Picea*-*Fagus* (mixed; n=6) and *Fagus*-*Fagu*s (mono; n=6). All plants were grown outside at Freiburg university campus (48°00’49.1"N 7°49’59.1"E, Germany) for one year until the beginning of experiment and rotated every two weeks to avoid potential effects of microclimates on plant development. In May 2022, plants were transferred to two walk-in climate chambers (Thermotec, Weilburg, Germany) and acclimatized to a relative humidity of 60 %, air temperatures of 23°C/20°C (day/night), a day length of 13 h and 900 μmol m^2^ s^-1^ photosynthetic photon flux density (PPFD) at canopy level. Plants were regularly irrigated to maintain a volumetric water content of ∼20 %.

**Figure 1:**
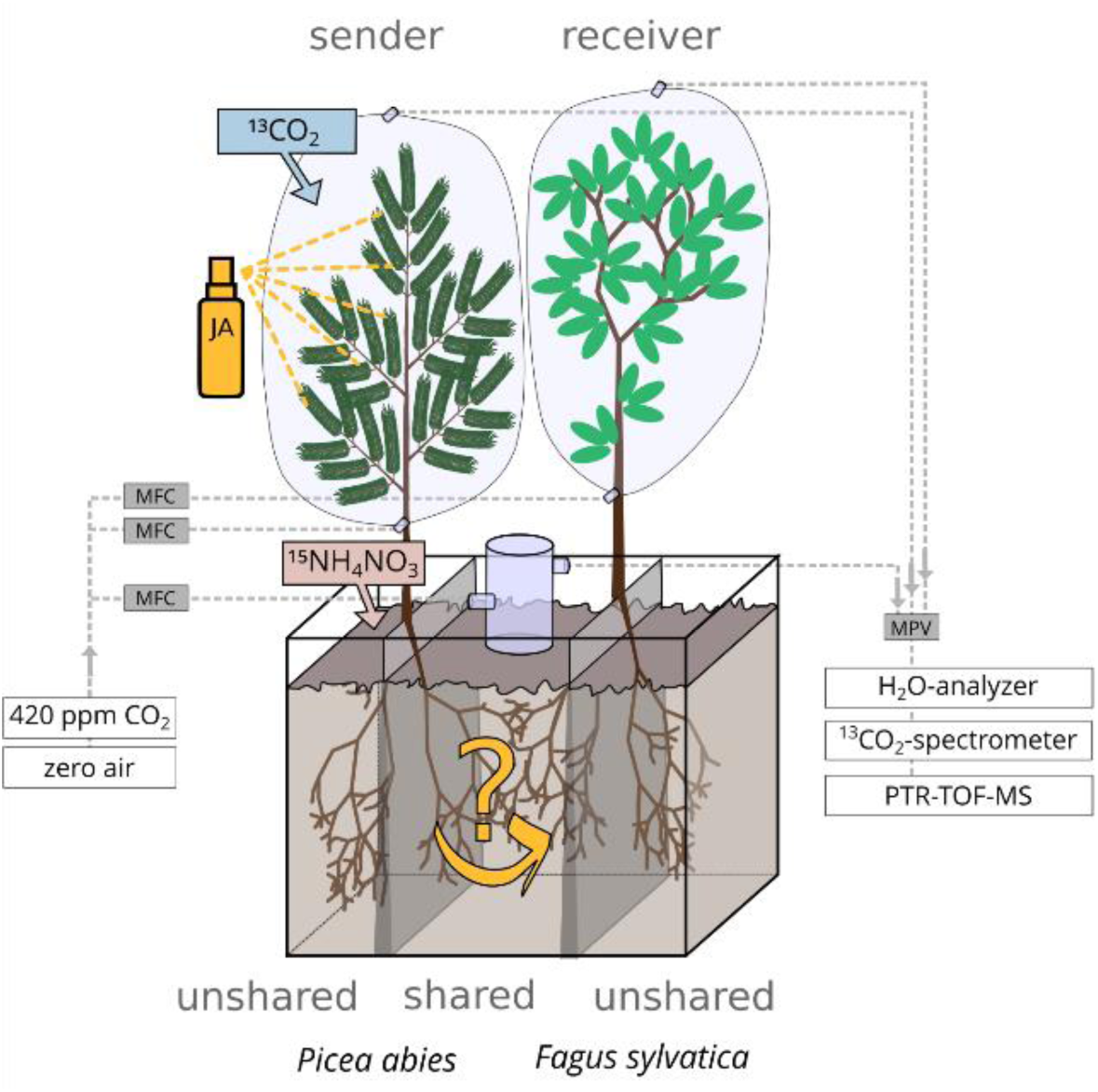
Split-root design to analyze belowground inter- and intraspecific interactions between *P. abies* and *F. sylvatica* (mixed, n = 6, depicted) and between two individuals of *F. sylvatica* (mono, n = 6). Jasmonic acid (JA) solution (10 mM) was applied to the shoot of sender plants to simulate herbivory. Additionally, sender plants were labeled with ^13^CO_2_ and ^15^NH_4_NO_3_ (100 mL 0.01mM) the day after JA treatment. Plants were shielded aboveground from each other to allow biotic interactions solely via the root system in the shared soil compartment. Throughout the experimental period of ten days, VOC emissions, ^13^CO_2_ and H_2_O fluxes from the shoots and soil surface were monitored with a custom build automated flow-through gas system installed in two walk-in climate chambers. VOC emissions were analyzed using a proton-transfer time-of-flight mass spectrometer (PTR-TOF-MS). MFC = mass flow controller, MPV = multi position valve.

### Continuous gas exchange and VOC measurements

Gas exchange and VOC emissions of shoots and soils were measured continuously over a period of ten days using an automated flow-through gas system installed in the two walk-in climate chambers. The system has been described in detail by Werner *et al*. (2020) and Meischner *et al*. (2024). In brief, it consists of a custom- made zero air generator; 14 in- and outlets for connecting plant enclosures or soil chambers; and an analyzer unit, as described below. For this experiment, the entire shoots of sender and receiver plants were enclosed in inert bags made of Nalophan foil, which served two purposes: firstly, to exclude airborne shoot-shoot signaling, and secondly, to analyze gas exchange and VOC emissions from the shoots. To measure soil gases, the shared soil compartment was equipped with an open-bottom chamber made of borosilicate glass (46.5 cm^2^ soil surface, 745 cm^3^ volume), which was inserted 3 cm into the soil three days before measurements were stared. Both, shoot enclosures and soil chambers, were flushed continuously with air (3 and 0.45 L min^-1^, respectively) which was adjusted to 400 ppm CO_2_ and purged of VOCs using an active charcoal filter and a platinum catalyst. The incoming and outgoing airstreams were analyzed in real-time for ^13^CO_2_, H_2_O, and VOC concentrations using an infrared laser spectrometer (Delta Ray IRIS, Thermo Fisher Scientific), a non- dispersive infrared gas analyzer (LI-850, LI-COR Environmental, Bad Homburg, Germany), and a proton- transfer-reaction time-of-flight mass spectrometer (PTR-TOF-MS 4000 ultra, Ionicon Analytic, Innsbruck, Austria), respectively. Two empty enclosures (blanks, one per climate chamber) were used to monitor the incoming airstream. A multi-position valve (VICI-Valco, Houston, TX, USA) was used to switch every 6 minutes between the 14 positions of the measurement system and the connected instruments had a minimum measurement interval of 20 seconds. The described setup allowed for the simultaneous analysis of four pairs of plants, resulting in three runs to measure all 12 pairs (May 8 – June 17, 2022).

### Experimental procedure

The experiment started with two days of control measurements before any treatment was applied (days -2 to - 1). On day zero, herbivory was simulated on the shoot of sender plants by the exogenous application of JA to induce a defense response. The following day (day 1), the same plants were labeled with ^13^CO_2_ and ^15^N- ammonium nitrate (^15^NH_4_NO_3_) in order to investigate the carbon allocation in stressed sender plants and the nutrient transfer between sender and receiver plants. JA application and stable isotope labeling are described in detail below. Gas exchange and VOC emissions of both, the sender and receiver plants, were monitored until day 7 after JA application. After completion of the continuous measurements, plants and soils were destructively sampled to analyze tissue and soil δ^13^C and δ^15^N (day 8). The sampled plant tissues included leaves/needles, branches, stems, coarse roots, fine roots, and, additionally, samples of rhizosphere soil were taken.

#### Jasmonic acid application

Exogenous application of JA was used to simulate herbivory on the shoots of sender plants. JA activates the JA signaling cascade (Wasternack & Hause, 2013). Inherently, it does not simulate the mechanical stress caused by insect feeding, which can result in the release of green leaf volatiles (GLVs) or VOCs from storage organs. Nevertheless, it induces VOC emissions similar to those induced by herbivores (Degenhardt & Lincoln, 2006; Li *et al*., 2019), while allowing highly standardized experiments (Waterman *et al*., 2019). In this study, approximately 1.5 mL of a 10 mM JA solution (JA dissolved in 10 vol.% ethanol and water, method adapted from Thaler *et al*., 1996) was sprayed evenly on the shoot of sender plants. For the treatment, plants were disconnected from the measurement system between 2:00 and 5:00 pm. Enclosures of the sender plants were opened to have access to the shoot whereas those of untreated receiver plants remained closed to prevent exposure to JA.

#### Pulse labeling with ^13^CO_2_

Sender plants were exposed for a period of four hours (1:30 – 5:30 pm) to 10% ^13^CO_2_, at a total CO_2_ concentration of 400 ppm. The desired concentration was obtained by adding 0.05 L min^-1^ of 99 % ^13^CO_2_ at a concentration of 400 ppm CO_2_ in nitrogen to 0.45 L min^-1^ of air with natural abundance of ^13^CO_2_ at the same CO2 concentration. During the pulse labeling, measurements were paused to avoid an ^13^CO_2_ oversaturation of the infrared spectrometer and the flow through the enclosures was reduced to 0.5 L min^-1^ for an efficient uptake of the ^13^C-label.

#### Labeling with ^15^NH_4_NO_3_

Plants were labeled with ^15^N by preparing a 0.1 mM solution of ^15^NH_4_NO_3_ (ammonium ^15^N, 98 %) in water, and adding 100 mL of this solution to the unshared compartment of sender plants during the ^13^CO_2_ labeling period (at 2 pm). To rinse the ^15^N solution into the soil, additional 50 mL of pure water was added.

### Analysis of tissue and soil δ^13^C and δ^15^N

For δ^13^C and δ^15^N analysis, samples were dried for 48 h at 60°C, pulverized and 2-5 mg of plant powder or 8 mg of soil powder were transferred into tin capsules. Samples were analyzed using an elemental analyzer (EA) (Vario Isotope Cube, Elementar, Langenselbold, Germany) coupled to an isotope ratio mass spectrometer (IRMS) (IsoPrime, Elementar, Langenselbold, Germany) as detailed in Werner *et al*. (2009) (see Methods S1 for further details). EA-IRMS data were referenced to IAEA-600 (caffeine) (δ^13^C = –27.8 ± 0.07) and δ ^15^N = 1.0 ± 0.16). The isotope ratios of the samples are expressed against international standards as δ (‰) notation according to equation (1):

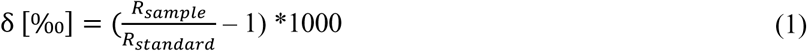

where R_sample_ and R_standard_ are isotopic ratios of a sample and the international standards (Vienna-Pee Dee Belemnite for carbon isotopes and atmospheric N₂ for nitrogen isotopes), respectively.

### Data acquisition and processing

VOC data were recorded with the PTR-TOF-MS operated in H_3_O^+^ ionization mode and controlled by IoniTOF software (version 4.4.69, Ionicon, Innsbruck, Austria). Raw data were processed with IDA software (version 2.2.0.7, Ionicon, Innsbruck, Austria) and calibrated either directly by measuring a multi-component gas mixture (Apel Riemer Environmental, USA) or using the quantification module in IDA software, if the compound was not present in the calibration mixture (see Methods S2 for further operation conditions and calibration details). For further data processing and statistical analyses, the software R (version 4.2.1, R Core Team, 2021) was used.

First, the average of each six-minute measurement was calculated, excluding the first two minutes to avoid transition effects from the previous measurement. Then, the background and limit of detection (LOD) were determined from blanks, with LOD defined as *x̅* + 2σ, being *x̅* the mean and σ the standard deviation of blank measurements. Values below LOD were set to zero and the background was subtracted from shoot and soil measurements before proceeding with the calculations of VOC fluxes. All formulas used for the calculation of VOC emission rates (nmol m ^−2^ h^−1^), assimilation rates A (µmol m^-2^ s^-1^), stomatal conductance for water vapor Gs (mmol m^-2^ s^-1^) and isotopic ratios of CO_2_ from dark respiration of the shoot or soil respiration δ^13^C (‰) are provided in the supplement (Methods S3). Statistical analysis

To determine ^13^C or ^15^N enrichment in aboveground tissues (leaves/needles, branches, stems), pairwise comparisons of isotopic ratios were made between labeled sender plants and unlabeled receiver plants using one-sided (decreasing) paired t-tests (n = 5-6). Plant individuals served as the pairing variable. For belowground tissues (coarse roots and fine roots) and rhizosphere soil, three separate t-tests with the same specifications as described above were conducted: (a) comparing the unshared and shared soil compartments of sender plants, (b) comparing the sender and receiver plants within the shared soil compartment, and (c) comparing the unshared and shared soil compartments of receiver plants. By this, the propagation of ^13^C and ^15^N could be followed along a gradient from highly enriched sender plants to potentially weakly enriched receiver plants. The comparisons were conducted separately for hetero- and monospecific treatments.

To analyze the VOC emissions of receiver plants, mixed effect models were fitted for each compound with the function ‘lmer’ of the package ‘lme4’ (version 1.1-35.1; Bates *et al*., 2015), specifying the day since JA treatment as fixed factor and the plant individuals as random factor. Models were fitted separately for hetero- and monospecific treatments (mixed/mono). Before the models were fitted, data were aggregated by taking averages of the control phase, and each day after JA treatment (days 1-7) for each plant individual. Model assumptions (normal distribution of residuals and variance homogeneity) were checked by the Lilliefors and Levene tests and, if required, data were square-root-transformed. To obtain p-values of the fixed effects the package ‘lmerTest’ was used (version 3.1-3; Kuznetsova *et al*., 2017) and the coefficients of determination (r^2^) of the models was determined taking into account both fixed and random effects (conditional r^2^) with the function ‘r2’ of the package ‘performance’ (version 0.10.9; Lüdecke *et al*., 2021). Significance levels of p < 0.05 (∗), P < 0.01 (∗∗) and p < 0.001 (∗∗∗) were applied for all hypothesis tests.

## Results

### Nutrient transfer between sender and receiver plants

Fine roots of sender plants labeled with ¹⁵NH₄NO₃ were ¹⁵N-enriched by 13.7 ± 3.6 ‰ and 16.00 ± 4.9 ‰ in the mono- and heterospecific treatment respectively, relative to the roots of receiver plants in the unshared compartment. Coarse roots showed a similar pattern but approximately 3 ‰ less label uptake. As illustrated in Fig. 1A, ¹⁵N enrichment was higher in the shoots than in the roots of labeled plants: *P. abies* needles were enriched by 27 ± 2.6 ‰ and *F. sylvatica* leaves by 33.5 ± 3.3 ‰, compared to unlabeled receiver plants. ¹⁵N- labeled sender plants also allocated ¹⁵N to the roots growing in the shared soil compartment, though to a lesser extent, with a mean Δ¹⁵N of 1.9 ± 0.69 ‰ averaged across both root types and species combinations (Fig. 1A). Importantly, in receiver plants, coarse roots in the shared compartment were significantly enriched in ^15^N compared to roots in the unshared compartment, by 1.1 ± 0.5 ‰ (p < 0.05) and 0.7 ± 0.5 (p < 0.05) in the hetero- and monospecific treatment, respectively (Fig. 2 D). Thus, roots, which shared the soil with labeled sender plants were enriched in ^15^N, regardless of whether the sender plant was *P. abies* or *F. sylvatica* (Fig. 2 D). In the heterospecific treatment, also fine roots of receiver plants were significantly enriched in ^15^N (p < 0.05) while no such difference was detected in the monospecific treatment (Fig. 2 D). Consequently, there is evidence for active root-root interactions in both species combinations, leading to ^15^N transfer from sender to receiver plants, which was more pronounced with *P. abies* as a sender, compared to *F. sylvatica*.

**Figure 2:**
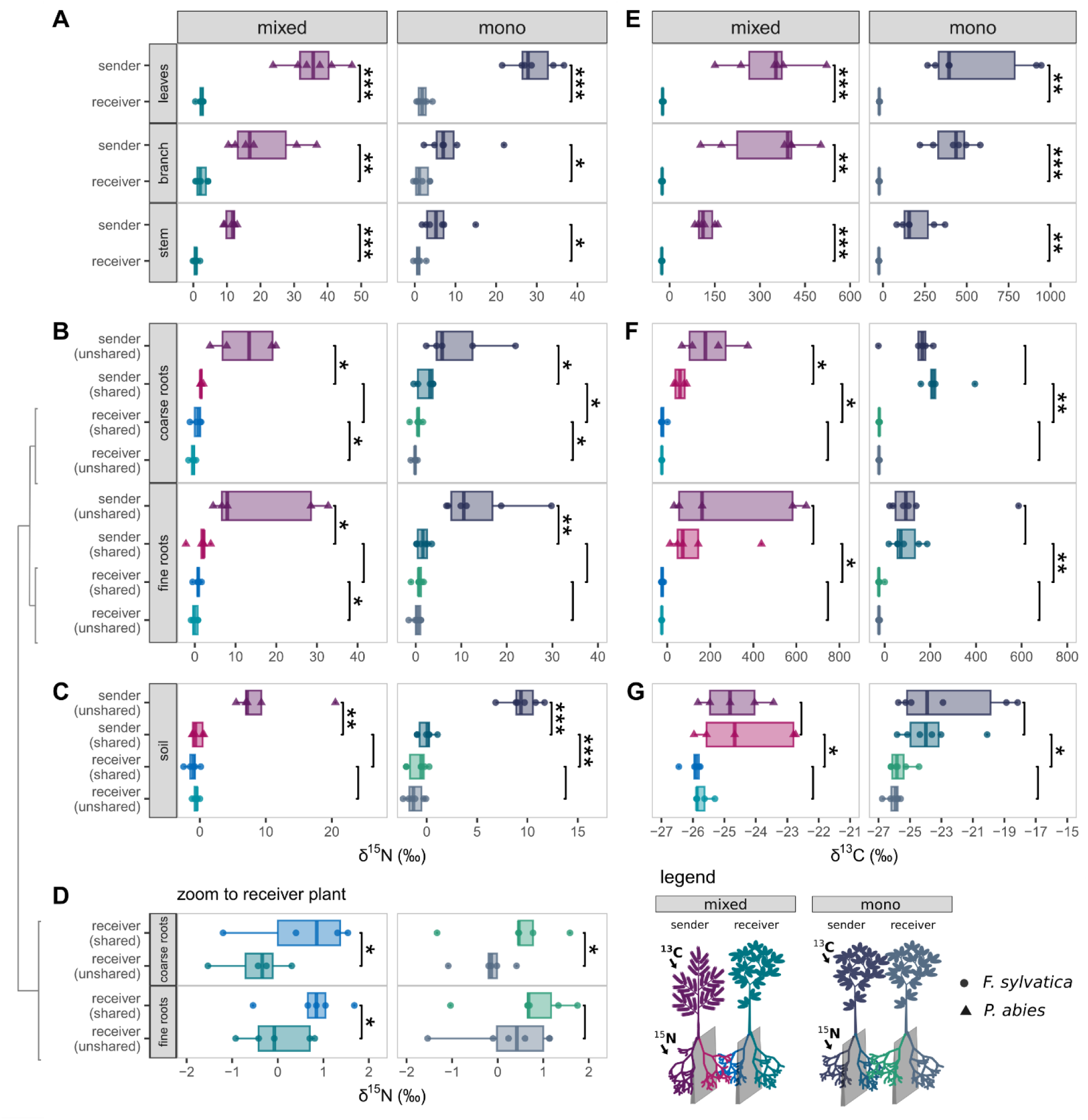
Nitrogen (A-D) and carbon (D-G) isotope composition of shoots (A & E), roots (B & D) and rhizosphere soil (C) of *Picea abies* and *Fagus sylvatica* saplings planted in pairs of two in a split-root design. Sender plants were labeled with ^15^NH_4_NO_3_ (100 mL 0.1mM, applied to the unshared soil compartment) and ^13^CO_2_ (exposure to 10% ^13^CO_2_ for 4 h) 8 days before harvesting. The experiment was repeated with hetero- (*Picea-Fagus*) and monospecific (*Fagus-Fagus*) species combinations (n=6, each). Differences between groups (indicated by brackets) were tested with paired t-tests using plant individuals as pairing variable. Significance levels of p < 0.05 (*), p < 0.01 (**) and p < 0.001 (***) were applied.

Exposure of the sender plants to ^13^CO_2_ resulted in δ^13^C values of 332.2 ± 51.9 ‰ and 537.9 ± 125.8 ‰ in needles and leaves of *P. abies* and *F. sylvatica*, respectively (Fig. 2 E). Furthermore, the isotopic composition of CO_2_ from dark respiration demonstrates a strong label incorporation particularly in the first night and the dilution of labeled carbon pools over time (Fig. 3 A). From the labeled shoot of *F. sylvatica*, ^13^C was distributed uniformly across both parts of the root system divided into the unshared and shared soil compartments (Fig. 2 F). In *P. abies*, there was a tendency for preferential allocation of assimilated carbon to the unshared compartment.

**Figure 3:**
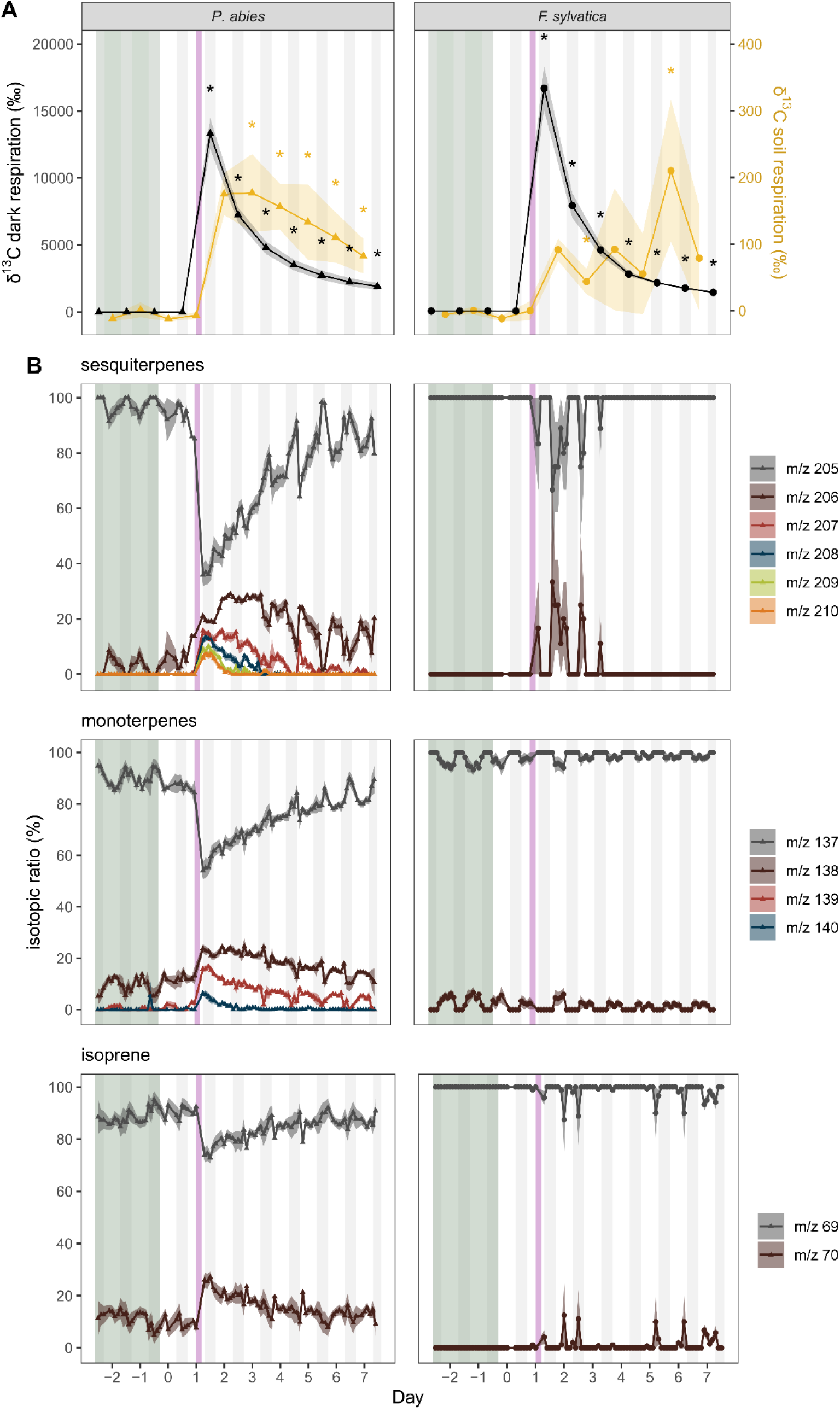
A: Carbon isotope composition of CO_2_ released by dark respiration from needles/leaves (black line) or soil respiration (orange line). Mean isotope composition per night for dark respiration and per day for soil respiration and standard errors are shown. Significant differences in isotopic rations (p < 0.05) compared to the control phase (day -1) are indicated by asterisks, as determined by Wilcoxon signed-rank tests. B: Ratios of unlabeled (gray) and labeled (colored) sesquiterpenes, monoterpenes, and isoprene. Mean isotope ratios and standard errors for 144-minute time intervals (≙10 values per day) are presented (n=6).

Hence, coarse roots in this compartment were significantly stronger enriched in ^13^C, compared to those in the shared compartment (δ^13^C = 215.3 ± 154.3 compared to 186.8 ± 48.5 ‰, p <0.05). However, these differences were limited to coarse roots and fine roots did not show a significant difference between compartments (Fig. 2 F). In the heterospecific treatment, δ^13^C ratios of respired CO_2_ had already reached their maximum the day after label application (Fig. 3 A), suggesting that assimilates are rapidly transported from shoots to roots. Conversely, in the monospecific treatment, there was slower carbon allocation to the roots by *F. sylvatica* compared to *P. abies* and δ^13^C ratios of respired CO_2_ reached their maximum on day 5. Interestingly, for both species combinations, rhizosphere soil of labeled plants in the shared compartment was slightly, but significantly (p < 0.05), enriched in ^13^C (δ^13^C = –24.5 ± 0.6 and –23.7 ± 0.8 ‰ in the hetero- and monospecific treatment, respectively) compared to rhizosphere soil of unlabeled plants in the same compartment (δ^13^C = -25.9 ± 0.1 and -25.6 ± 0.3 ‰ in the hetero- and monospecific treatment, respectively) (Fig. 2 G) indicating the exudation of organic molecules from the roots of sender plants into the soil.

### Response of sender plants and soil VOCs to simulated herbivory

*P. abies* and *F. sylvatica* exhibited different responses to simulated herbivory. In *P. abies* shoots, isoprenoid, benzene and MVK emissions increased sharply in response to JA treatment (Fig. 4). Conversely, acetone emissions decreased (Fig. 4). Notably, there was a strong decrease in the proportion of unlabeled sesquiterpenes, monoterpenes and isoprene with a rapid incorporation of up to five ^13^C atoms in sesquiterpenes (Fig. 3 B). Furthermore, for sesquiterpenes, the rates of ^13^C incorporation varied diurnally, with higher incorporation rates observed during the day compared to the night (Fig. 3 B), suggesting that different carbon sources were used for their biosynthesis depending on the time of day.

**Figure 4:**
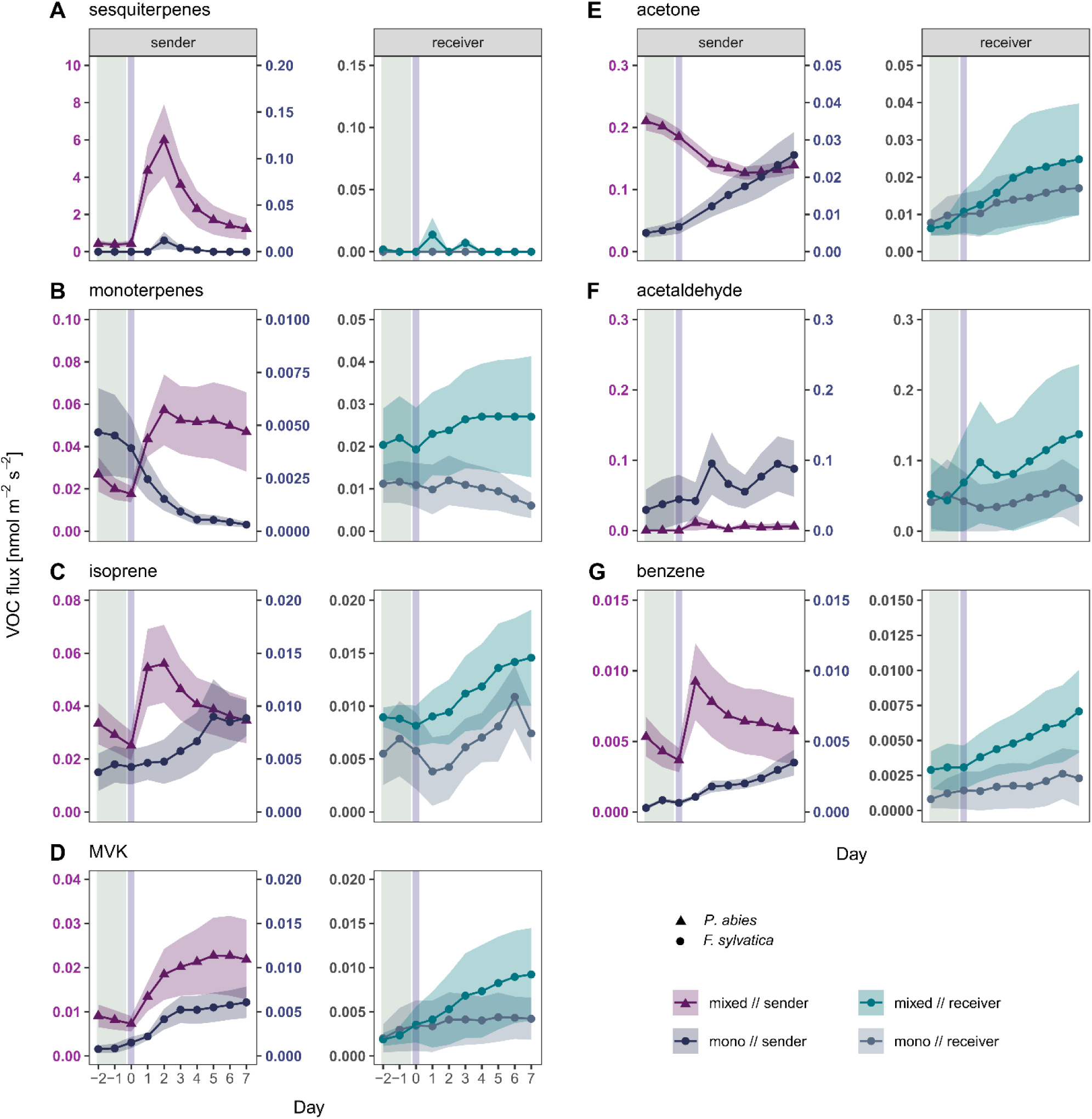
Daily averages and standard errors (n=6) of VOC emissions from shoots of *Picea abies* (triangles) and *Fagus sylvatica* (circles) pairs planted in a split-root design to investigate root-root interactions. Shoots of sender plants were treated with 10 mM jasmonic acid solution on day zero, receiver plants remained untreated. The shoots of sender and receiver plants were shielded from each other to avoid aboveground signaling. The green background indicates the control phase before JA application. VOC emissions were measured continuously during ten days by PTR-TOF-MS. Note different scales for VOC fluxes from *P. abies* and *F. sylvatica*. MVK = methyl vinyl ketone. For statistics see Table 1.

**Table 1:**
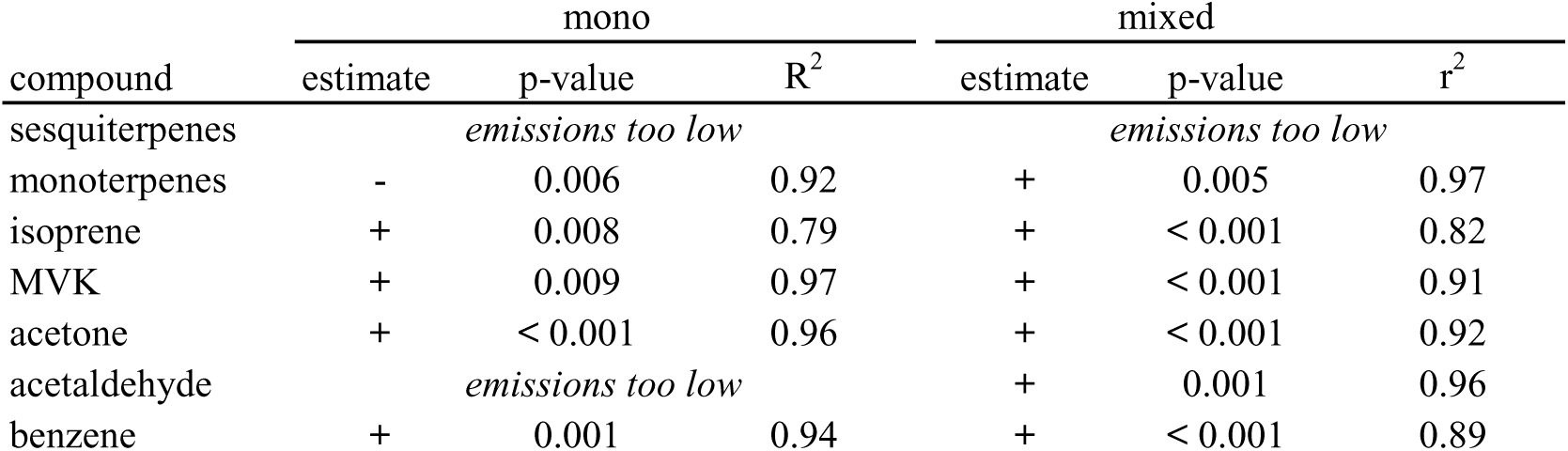
Mixed effect models of VOC emissions from receiver plants days after with JA treatment as fixed effect and plant individuals as random effects. Models were fitted separately for both species combinations (mixed/mono). Data of acetaldehyde (mono), benzene (mono) and acetone (mixed) were square root transformed to meet the model assumptions (normal distribution of residuals and variance homogeneity).

Despite the fact that total VOC emissions from *F. sylvatica* were by several orders of magnitude lower compared to *P. abies*, they also differed compared to constitutive emissions in response to simulated herbivory (Fig. 4): emissions of MVK, acetone, acetaldehyde and benzene increased, while monoterpene emissions decreased. Sesquiterpene emissions were low and close to the LOD. Furthermore, there was no incorporation of ^13^C label into monoterpenes (Fig. 3 B). Even though sesquiterpenes were only emitted in low amounts, they exhibited strong ^13^C incorporation from freshly assimilated carbon. (Fig. 3 B). In both species, there was a slight reduction in net carbon assimilation and a significant increase of dark respiration rates on the days following direct JA treatment (Fig. S1).

Based on their different VOC profiles, *P. abies* can be characterized as a species with high emission rates and strong responses to the JA treatment and, while *F. sylvatica* had lower VOC emission rates and reacted to a lower extent to JA treatment.

### Response of receiver plants to simulated herbivory on shoots of neighboring plants

Importantly, unstressed receivers (*F. sylvatica*) growing next to JA-stressed con- and heterospecific neighbors exhibited induced VOC emissions after JA application on their neighbors’ shoots with a time delay of approximately one day (Fig. 4). Specifically, receivers significantly increased monoterpene (p = 0.005), isoprene (p < 0.001), MVK (p < 0.001), acetone (p < 0.001), acetaldehyde (p = 0.001) and benzene emissions (p < 0.001) (Table 1), when the sender was *P. abies*. Interestingly, these emissions showed a high degree of similarity with those of directly treated individuals of *F. sylvatica* (Fig. 4). When receivers were situated next to a stressed conspecific (*F. sylvatica*), there was a significant increase in the emission, isoprene (p = 0.008), MVK (p = 0.009), acetone (p < 0.001) and benzene emissions (p = 0.001) following the JA treatment on senders, while monoterpene emissions decreased (p = 0.006). Overall, the response of *F. sylvatica* receivers was more pronounced, when they were located next to *P. abies* compared to a conspecific (Fig. 4). Noteworthy, receiver plants had similar constitutive VOC emission before JA application to sender plants independent of the species combination and the differences in VOC emissions only emerged after JA treatment. Thus, the species combination modulated the response intensity of the receiver plants. In contrast to sender plants, which have been directly treated with JA, the photosynthetic gas exchange, i.e. assimilation and dark respiration rates, remained at pre-stress levels in the receiver plants (Fig. S1).

### Soil VOC emissions

Finally, the pattern of *F. sylvatica* as a weak sender and *P. abies* as a strong sender was also reflected in soil VOC emissions. In both species combinations, the composition of soil VOCs consisted primarily of formic acid, with smaller amounts of carbonyls (acetaldehyde and acetone) and toluene. In the heterospecific treatment, methyl salicylate, was also emitted from the soil, and formic acid emissions increased significantly (p < 0.05) on the day following JA treatment, remaining elevated until the end of the measurement period. In the monospecific treatment, formic acid emissions from soils also increased, albeit to a lesser extent.

These data demonstrate that, aboveground JA treatment had a profound impact on both aboveground VOC emissions of sender and receiver plants as well as VOC emissions from the soil, especially with *P. abies* as sender plant.

## Discussion

We investigated belowground signaling in a split root-design between *P. abies* and *F. sylvatica*, two dominant tree species in temperate forests. We show that *F. sylvatica* responds to belowground stress cues triggered by JA pathway activation in neighboring plants with increased shoot emissions of oxygenated VOCs and benzenoids. This response reflects a complex signaling cascade – from the shoots to the roots of sender plants, and then, potentially via the roots and soil, to the shoots of the receiver plants (Fig. 6). Notably, the response of receiver plants was more pronounced when the sender plant was a strong emitter as *P. abies* than when it was a conspecific *F. sylvatica*, characterized as a weaker sender. This suggests that the signal intensity determines the magnitude of the receiver’s response. Belowground interactions were further confirmed by transfer of ^15^N from senders to receivers and the exudation of ^13^C into the soil.

**Figure 5:**
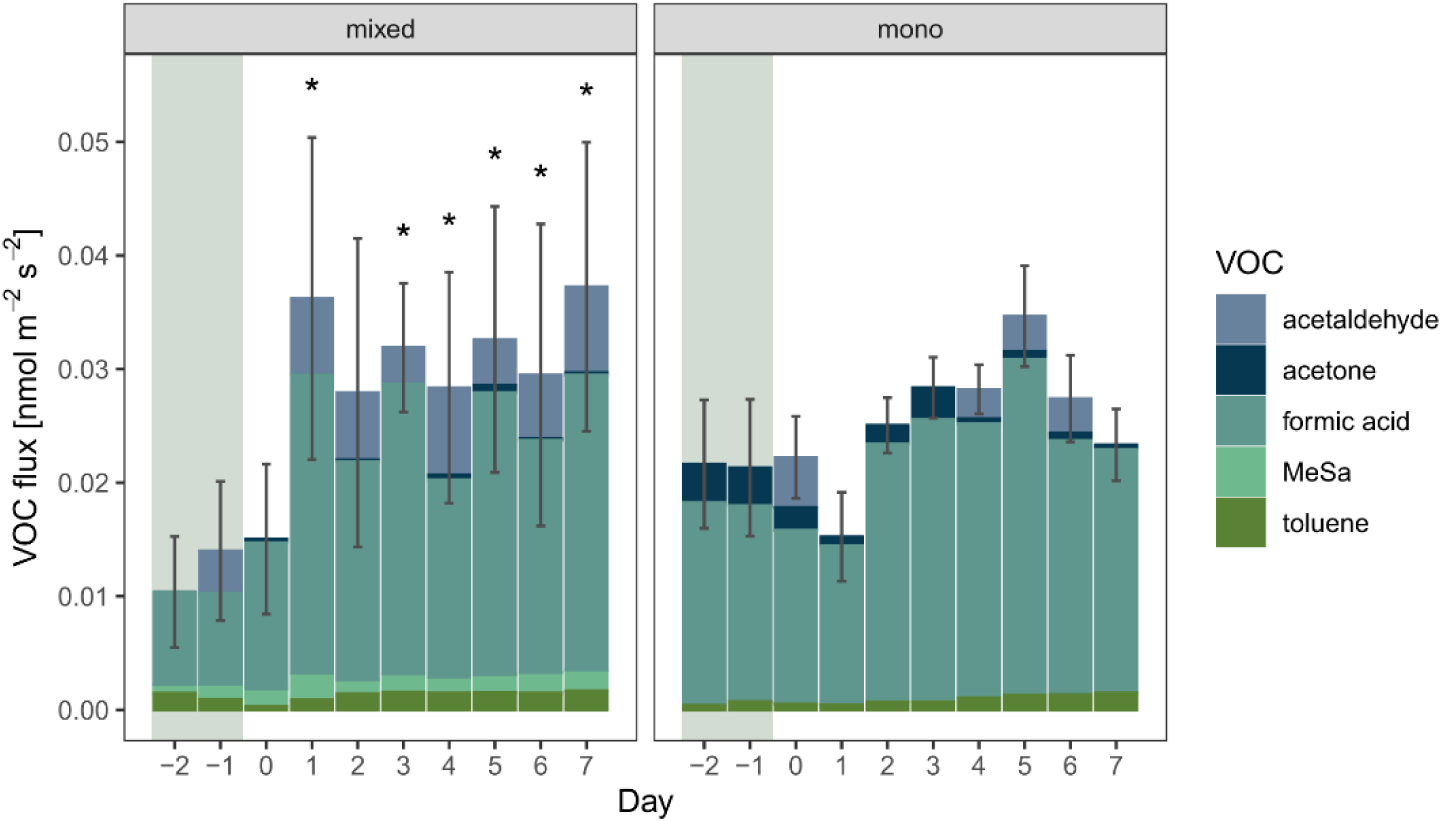
Soil VOC emissions from soil compartments shared by either a *Pica abies* and a *Fagus sylvatica* sapling (mixed, n = 6) or two *F. sylvatica* saplings (mono, n = 6). Open-bottom chambers connected to an automated measuring system were used for measuring VOCs by PTR-TOF-MS. Mean daytime VOC emissions per day and standard errors of the total soil VOC emissions are shown. Asterisks indicate significant differences in formic acid emissions compared to the control phase (day -1) and were determined using Wilcoxon signed-rank tests. MeSa = methyl salicylate.

**Figure 6:**
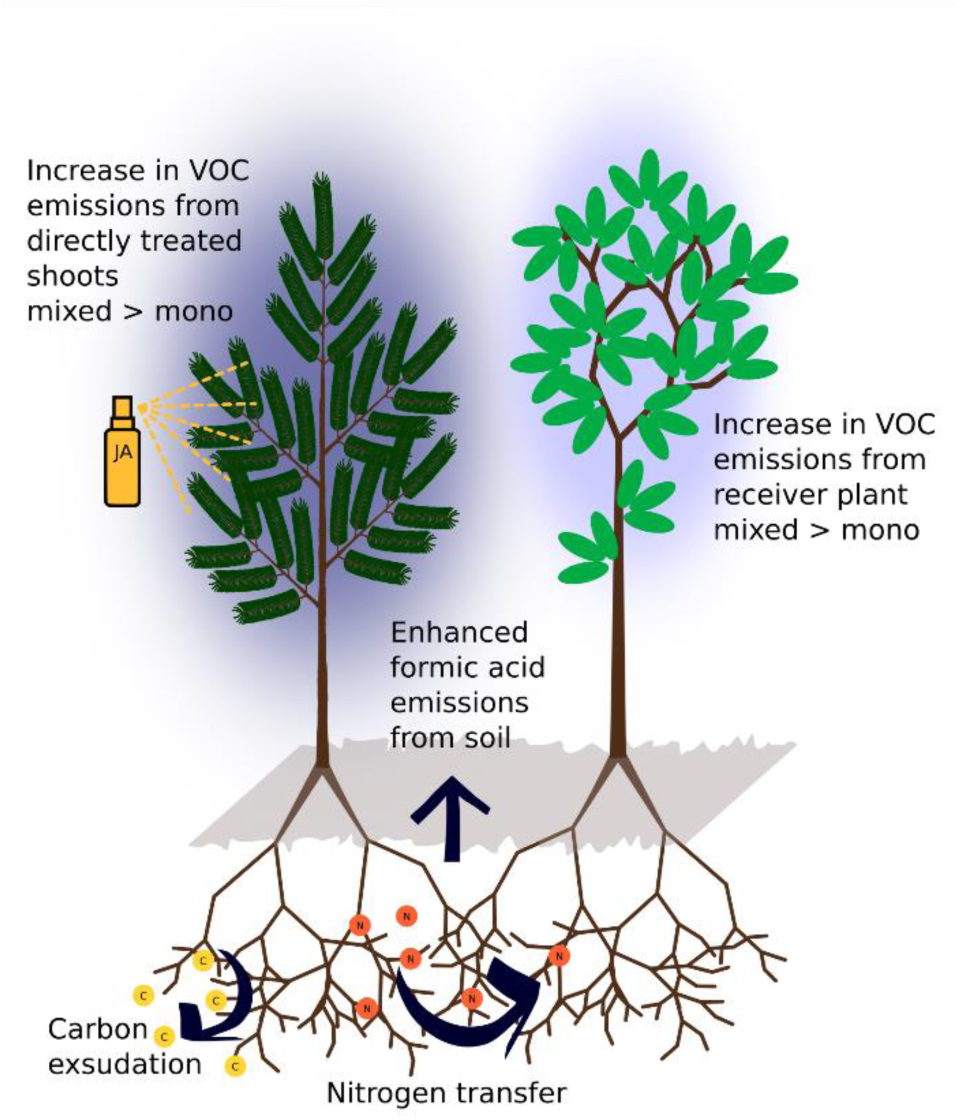
Summary of split-root experiment conducted with pairs of *P. abies* and *F. sylvatica* saplings (mixed, n = 6, depicted), or two *F. sylvatica* saplings (mono, n = 6). Shoots were shielded from each other to exclude airborne VOC signaling.

### Nitrogen transfer and carbon exudation

Nitrogen translocation between trees has been documented in various plant species and is frequently studied in the context of mycorrhiza-mediated tree-tree interactions (He *et al*., 2003, 2019; Fellbaum *et al*., 2014; Teste *et al*., 2015). Both *F. sylvatica* and *P. abies* form ectomycorrhizal associations, including shared fungal partners such as *Paxillus involutus* and *Russula ochroleuca* (Schirkonyer *et al*., 2013; Rosinger *et al*., 2018).

Also, N uptake by *F. sylvatica* from *Paxillus involutus* has been previously demonstrated (Finlay *et al*., 1989), making ectomycorrhizal networks a plausible transfer route for nitrogen. However the role of common mycorrhizal networks in direct plant-to-plant nutrient exchange remains controversial in recent literature (Karst *et al*., 2023; Klein *et al*., 2023; Robinson *et al*., 2024). Alterative or additional transfer pathways include the uptake of N from nitrogenous root exudates (e.g. amino acids) or diffusion driven solution of N in the soil water and subsequent uptake by receiver plants (Biernath *et al*., 2008). In our experiment, the difference in δ^15^N between shared and unshared soil compartment in the receiver plants’ coarse roots was slightly greater for *F. sylvatica* paired with *P. abies* than with a conspecific (Δ δ^15^N 1.1 ± 0.5 vs. 0.7 ± 0.5 ‰, respectively), suggesting greater N transfer from *P. abies* than from *F. sylvatica*. However, this may reflect species-specific N acquisition strategies, as *P. abies* exhibited generally a higher ^15^N enrichment in the shoot than *F. sylvatica* (Fig. 2 A) consistent with previously reported differences in N uptake mode of both species (Gessler *et al*., 1998).

The enrichment of ^13^C in the rhizosphere of sender plants after aboveground ^13^C fumigation indicates that non- volatile metabolites were exuded into the soil in considerable amounts. Under well-watered conditions, the proportion of net carbon assimilation allocated to total exudation typically ranges between 0.1% and 0.2% for both tree species examined in this study (Brunn *et al*., 2022). Root exudates are chemically highly diverse and their exudation rates have been shown to be responsive to foliar herbivory (Rolfe *et al*., 2019; Xing *et al*., 2024). Moreover, root exudates play a critical role in belowground interactions (Wang *et al*., 2021; Mathieu *et al*., 2024) and may have contributed to the transfer of signals from sender to receiver plants in the present study. Notably, root exudates and root-emitted volatiles can function directly as chemical cues (e.g. ethylene), or induce indirect effects on neighboring plants by modulating soil microbial communities and their activity (Hu *et al*., 2018, 2025; Kong *et al*., 2021). An indicator of such microbial modulation in this study is the observed increase in formic acid emissions from soils following aboveground JA application, which was significant in the heterospecific, but not in the monospecific treatment (Fig. 5). Formic acid is a low molecular weight organic acid commonly emitted from soils with emission rates strongly influenced by soil moisture and temperature (Sanhueza & Andreae, 1991; Mielnik *et al*., 2018; Meischner *et al*., 2022). In a previous study on *P. abies*, no formic acid emissions were detected from soil-free roots (Meischner *et al*., 2024), supporting a microbial origin of this compound. To preserve the delicate root-to-soil and root-to-root interfaces, roots were not excavated for direct VOC measurements in this study. Instead, net VOC emissions were recorded at the soil surface, representing the net emissions from root and microbial production and microbial and abiotic VOC uptake (Rinnan & Albers, 2020; Jiao *et al*., 2023). Thus, these findings suggest that the aboveground application of JA influenced belowground soil chemistry and microbial activity in the rhizosphere, and ^15^N signatures indicate active root-root interactions between sender and receiver plants.

### Stronger reaction of Picea abies to simulated herbivory compared to Fagus sylvatica

Among the directly JA-treated sender plants, *P. abies* exhibited a stronger increase in total VOC emissions, particularly of isoprenoids (isoprene, monoterpene and sesquiterpenes), exceeding those of *F. sylvatica* by several orders of magnitude (Fig. 4). Monoterpenoids and sesquiterpenoids are well known for their defensive functions, including herbivore deterrence and their roles as signaling molecules in biotic interactions, amongst others (Gershenzon & Dudareva, 2007; Unsicker *et al*., 2009; Ditengou *et al*., 2015). Notably, exposure of tomato (*Solanum lycopersicum*) shoots to ß-caryophyllene, a sesquiterpene also present in the VOC blend emitted from *P. abies* needles and roots (Daber *et al*., 2025), has recently been shown to elicit salicyclic acid emission from roots, promoting the growths of the beneficial rhizobacterium *Bacillus amyloliquefaciens* in the rhizosphere (Kong *et al*., 2021). This illustrates the significance of shoot-root communication in plant defense and highlights the influence of aboveground signals on rhizosphere microbial communities.

In contrast, *F. sylvatica* emitted lower amounts of sesquiterpenes and even showed a decrease in monoterpene emissions following JA treatment with low rates of ^13^CO_2_ incorporation. Instead, JA treatment led to increased emission of oxygenated VOCs, including acetaldehyde, acetone and MVK, as well as benzene.

Benzene has been shown to be synthesized by plants via the shikimate pathway and is thought to function as a signaling molecule in plant defense (Misztal *et al*., 2015), whereas MVK is an oxidation product of isoprene (Loreto & Schnitzler, 2010) or originates from isoprene hydroxyperoxides (Canaval *et al*., 2020). Yet, the role of acetaldehyde and acetone in the stress response of plants is less well understood. Both compounds are closely linked to primary metabolic processes: acetaldehyde is a decarboxylation product of pyruvate (Kreuzwieser *et al*., 1999, 2000; Graus *et al*., 2004; Loreto & Schnitzler, 2010), while acetone is potentially synthesized from decarboxylated pyruvate in the cytosol (Jardine *et al*., 2010).

### Increased volatile organic compound emissions in receiver plants are dependent on sender species

Notably, emissions of acetaldehyde, acetone, MVK, and benzene, which were sensitive to direct JA application to *F. sylvatica*, also began to rise with a one-day delay in neighboring receiver plants. This effect was more pronounced when the neighboring plant was a strong sender as *P. abies*. *F. sylvatica* evoked similar, but weaker responses in receiver plants and can, thus, be characterized as weak sender. Intraspecific signal transfer has been previously observed between *Pseudotsuga menziesii* and *Pinus ponderosae,* where both, mechanical damage and insect defoliation of *P. menziesii,* increased the activity of the defense enzymes peroxidase, polyphenol oxidase and superoxide dismutase in *P. ponderosae* (Song *et al*., 2015). However, most research on belowground propagation of stress signals has focused on herbaceous species (Rasheed *et al*., 2023). For instance, elevated methyl salicylate emissions from bean (*Vicia faba*) plants were triggered by aphid feeding in both infested plants and their uninfested neighbors, provided their roots were not separated by a mesh impermeable to mycorrhizal fungi (Babikova *et al*., 2013b). Notably, the response in receiver plants began within 24 hours (Babikova *et al*., 2013a), similar to the time lag observed in our study. Song *et al*. (2010) also emphasized the importance of mycorrhizal fungi in the belowground plant-to-plant signaling and suggested that the JA signaling pathway is activated in receiver plants adjacent to conspecific neighbors infested with *Spodoptera litura* caterpillars. This aligns with our observation that similar VOCs were induced in both JA-treated sender plants and their neighbors, supporting the idea of shared signaling pathways. Although experiments that are using root meshes with varying pore sizes provide valuable insights into the involvement of mycorrhizal networks in belowground signaling (e.g. in Song *et al*., 2010; Babikova *et al*., 2013b), more experiments are needed that exclude aboveground signaling to further investigate belowground communication. As with airborne signaling, where mechanisms of VOC perception are currently a major focus of research (Loreto & D’Auria, 2022; Kessler *et al*., 2023; Bergman *et al*., 2025), many fundamental questions regarding belowground signaling and perception require further investigation.

In conclusion, this work provides evidence for a complex signal cascade between two dominant temperate tree species that is mediated solely via the soil, since airborne signals were excluded (Fig. 6). Furthermore, we found that receiver plants responded more strongly when grown next to *P. abies*, a species characterized by strong VOC emissions, than to conspecific *F. sylvatica* saplings. This finding could imply that *F. sylvatica* growing in heterospecific stands with *P. abies* could benefit from its ’communicative’ neighbor by eavesdropping on information about surrounding herbivory and increasing their defenses at an early stage of herbivory propagation.

## Author Contributions

MM, SH, JK, JPS and CW designed the study and discussed data. MM analyzed the data and wrote the manuscript with input from all authors.

## Acknowledgements

The authors would like to thank Eva Schottmüller for gardening the experimental plants, Monika Siegel and Niklas Wolf for their assistance with sample processing and preparation, Anne-Marie Schiphorst and Alexandra Paul for performing the EA-IRMS analyses, and Erik Daber and Michael Rienks for providing technical support with the automated measurement system. Erik Dabers’s assistance with the PTR-TOF-MS measurements is also gratefully acknowledged.

## Data availability

The data that support the findings of this study are available from the corresponding author upon reasonable request.

## Supporting information

Methods S1: EA-IRMS operating conditions

Methods S2: PTR-TOF-MS operating conditions and calibration details

Methods S3: Calculation of VOC fluxes and gas exchange parameters

Figure S1: Gas exchange parameters

