## Supporting information for "Eavesdropping roots: *Fagus sylvatica* detects belowground stress signals from conspecific and heterospecific (*Picea abies*) neighbors, triggering increased shoot VOC emissions"

### Eavesdropping roots: *Fagus sylvatica* perceives stress signals belowground from conspecific and non-conspecific (*Picea abies*) neighbours and responds with enhanced VOC emissions from the shoot

Mirjam Meischner[*1], Simon Haberstroh[1], Jürgen Kreuzwieser[1], Jörg-Peter Schnitzler[2], Christiane Werner[1]

[1] Ecosystem Physiology, University of Freiburg, 79110 Freiburg, Germany
[2] Research Unit Environmental Simulation, Helmholtz Zentrum München, 85764 Neuherberg, Germany

**The following supporting information is available for this article:**

Methods S1: EA-IRMS operating conditions
Methods S2: PTR-TOF-MS operating conditions and calibration details
Methods S3: Calculation of VOC fluxes and gas exchange parameters
Figure S1: Gas exchange parameters

#### Methods S1: EA-IRMS operating conditions

EA Mode of Operation: CN
Trap Current $CO_2$: 100 µA
Trap Current $N_2$: 600 µA
Dilution $CO_2$: 6,5% $CO_2$
Dilution $N_2$: none - 100% $N_2$

Table S1: Sample weight for EA-IRMS analysis [mg]

| | *Picea abies* | | *Fagus sylvatica* | |
|---|---|---|---|---|
| | sender | receiver | sender | receiver |
| leaf | 3 | 3 | 3 | 2 |
| branch | 2 | 2 | 3 | 3 |
| stem | 5 | 5 | 5 | 5 |
| soil | 8 | 8 | 8 | 8 |
| fine root | 2 | 2 | 3 | 3 |
| coarse root | 2 | 2 | 2 | 2.2 |

**Methods S2: PTR-TOF-MS operating conditions and calibration details**

Ionization mode: $H_3O^+$
Drift tube temperature: 80°C
Drift pressure of 2.7 mbar
Drift voltage of 556 V
E/N*: 107 Td

*electric field strength/number density of the drift tube buffer gas molecules

Table S2: Compounds present in the multi-component gas mixture (Apel Riemer Environmental, USA) used for calibration of VOC data and the sensitivity of the PTR-TOF-MS for these compounds.

| **Compound** | **Protonated Mass** | **Sensitivity (cps/ppb)** |
|---|---|---|
| Formaldehyde | 31.018 | 6.10 |
| Methanol | 33.034 | 23.60 |
| Acetonitrile | 42.034 | 65.40 |
| Acetaldehyde | 45.034 | 58.50 |
| Acetone | 59.049 | 72.70 |
| Dimethyl Sulfide | 63.026 | 54.20 |
| Isoprene | 69.070 | 44.20 |
| Methacrolein+MVK | 71.049 | 55.00 |
| Methyl Ethyl Ketone | 73.068 | 68.60 |
| Methyl Acetate | 75.044 | 51.50 |
| Benzene | 79.054 | 47.60 |
| Toluene | 93.070 | 68.20 |
| 1-Hexanol | 103.112 | 0.20 |
| Benzaldehyde | 107.049 | 64.60 |
| 1,2,4-Trimethylbenzene | 121.101 | 37.80 |
| p-Cymene | 135.117 | 11.90 |
| Alpha-Pinene | 137.133 | 59.00 |
| 1,8-Cineol | 155.144 | 1.40 |
| Beta-Caryophyllene | 205.195 | 8.20 |

Sensitivity was obtained using a liquid calibration unit (Ionicon Analytic, Innsbruck, Austria). Compounds present in the calibration gas were exported as cps values from IDA software and calibrated directly by dividing the cps values by the sensitivity. The ppb values of compounds that were not contained in the calibration gas were determined using the quantification module in IDA software based on the saved transmission rates which were obtained using the calibration gas (Table S2), all quantification relevant instrumental parameters including drift tube voltage, pressure and temperature and reaction rate coefficients (k-rates) of the detected compounds.

**Methods S3: Calculation of VOC fluxes and gas exchange parameters**

VOC fluxes

To retrieve VOC fluxes and gas exchange parameters, the molar flow u (mol $s^{-1}$) through the cuvettes was determined with Equation (1)

$$u = \frac{V}{t} * \frac{p}{R * T} \tag{1}$$

where V is the gas volume ($m^3$), t is time (s), p is air pressure (Pa), R is the molar gas constant (J $mol^{-1}$ $K^{-1}$) and T is temperature (K).

To retrieve VOC fluxes e (mol $m^{-2}$ s–$^{1}$) Equation (2) was used:

$$e = \frac{u}{w} * (VOCo - VOCa) \tag{2}$$

where w is the leaf/needle area or soil surface ($m^2$) and VOCo and VOCa are the concentrations of VOCs (mol $mol^{-1}$) at the cuvette outlet and inlet, respectively.

Soil respiration rates

Analogously, soil respiration rate Rs (mol g-1 s-1) was calculated with Equation (3):

$$Rs = \frac{u}{w} * (r\_co - r\_ca) \tag{3}$$

where w is the soil surface ($m^2$) and r_co and r_ca are the concentrations of $CO_2$ (mol $mol^{-1}$) at the outlet and inlet of the open-bottom soil chamber, respectively.

Transpiration and assimilation rates

First the transpiration rate E (mol $m^{-2}$ $s^{-1}$) was determined according to von Caemmerer & Farquhar (1981) with Equation (4):

$$E = \frac{u}{s} * \frac{wo - wi}{1 - wo} \tag{4}$$

where s is leaf area ($m^2$) and wo and wi are the concentrations of $H_2O$ (mol $mol^{-1}$) at the outlet and inlet of the cuvette, respectively.

Finally, assimilation rate (mol $m^{-2}$ $s^{-1}$) was calculated using Equation (5):

$$A = \frac{u}{s} * \left(\frac{1 - wi}{1 - wo}\right) * (ci - co) - E * ci \tag{5}$$

being co and ci the concentrations of $CO_2$ (mol $mol^{-1}$) at the outlet and inlet of the cuvette, respectively

Stomatal conductance

The absolute humidity inside the cuvette $AH_c$ (g m$^{-3}$) based on the measured concentration of $H_2O$ (mol mol$^{-1}$) at the cuvette inlet was calculated with Equation (6):

$$AH_c = \frac{wi * m}{V_m} \tag{6}$$

where m is the molecular weight of water (= 18.01528 g mol$^{-1}$) and $V_m$ the molar volume of an ideal gas at standard temperature and pressure (22.4·10$^{-3}$ m$^3$ mol$^{-1}$).

The absolute humidity inside the leaf $AH_l$ was determined based on the measured temperature according to Steubing & Fangmeier (1992). First the saturation vapor pressure e (kPa) was calculated with equation (7):

$$P = 101{,}325 \exp(13{,}3185t - 1{,}976t^2 - 0{,}6445t^3 - 0{,}1229t^4) \tag{7}$$

where t is defined as $t = 1 - (Ts/T)$ and Ts as vapour temperature at standard air pressure (=373.15 K).

The vapor pressure was then converted into absolute humidity (g m$^{-3}$) using Equation (8) according to Lösch (2001):

$$AH_l = \left(\frac{2.17}{T}\right) * P * 1000 \tag{8}$$

The absolute water vapour pressure deficit (VPD) (g m$^{-3}$) is the difference between $AH_l$ and $AH_c$ (equation 9):

$$VPD = AH_l - AH_c \tag{9}$$

Finally, stomatal conductance for water vapour Gs (mol m$^{-2}$ s-$^{1}$) is calculated with Equation (10)

$$Gs = \frac{\frac{E * m}{VPD}}{Vm} \tag{10}$$

Isotopic ratios of $CO_2$ from dark respiration of the shoot or soil respiration

Isotopic rations $\delta^{13}C$ (‰) of $CO_2$ released by dark respiration of the shoot or soil respiration, was calculated based on the absolute differences of $CO_2$ and $^{13}C$ ratios at the inlet and outlet of the cuvette using equation (11)

$$\delta^{13}C\ (‰) = \frac{co * \delta^{13}CO_2 out - ci * \delta^{13}CO_2 in}{co - ci} \quad (11)$$

where $\delta^{13}CO_2$ *in* is $\delta^{13}CO_2$ at the inlet of the cuvette and $\delta^{13}CO_2$*out* at the outlet.

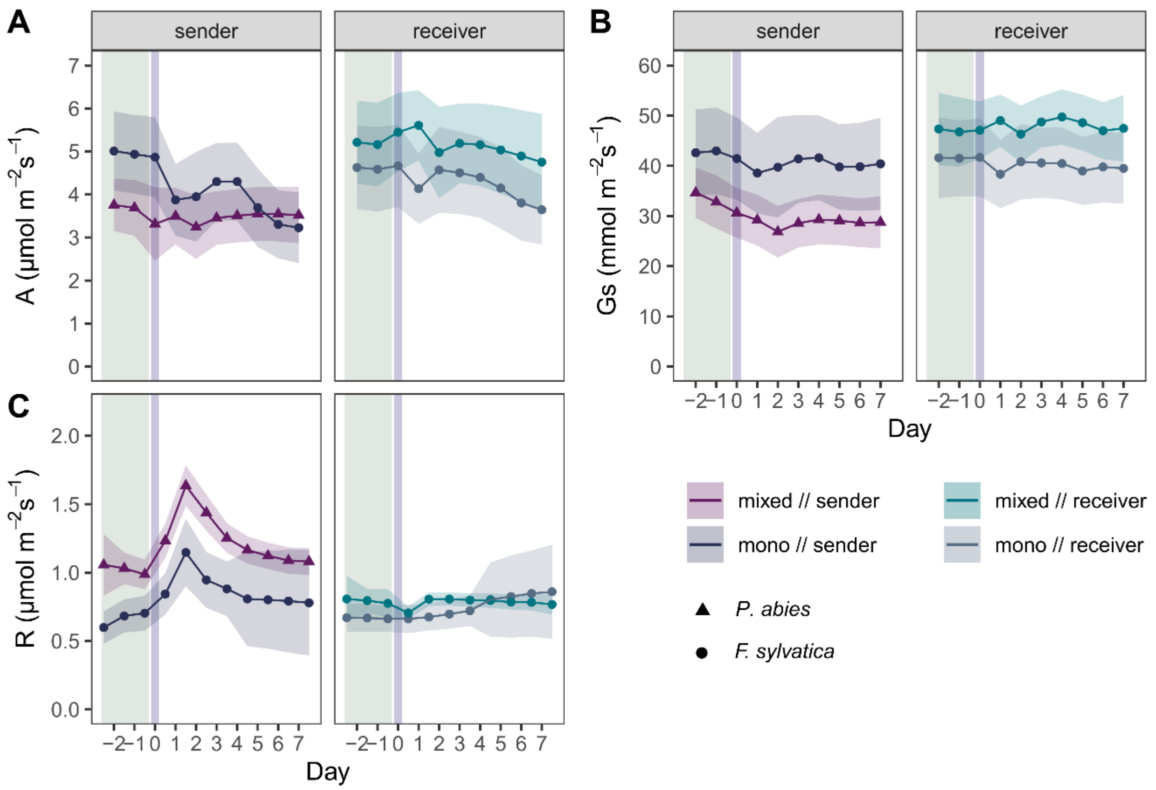


Figure S1: Daily means and standard errors of net assimilation rates (A) (panel A), stomatal conductance for H2O (Gs) (panel B) and dark respiration rates (R) (panel C) of *Picea abies* and *Fagus sylvatica* saplings (n=6). Trees were planted in pairs of two in the combinations *Picea-Fagus* (mixed) and *Fagus-Fagus* (mono) in a split-root design. Sender plants were treated with jasmonic acid (JA) solution on day zero (indicated by blue background) and receiver plants remained untreated. The green background indicates the control phase before JA treatment.
